# Generation of white-eyed *Daphnia magna* mutants lacking *scarlet* function

**DOI:** 10.1101/313395

**Authors:** Nur Izzatur Binti Ismail, Yasuhiko Kato, Tomoaki Matsuura, Hajime Watanabe

**Author notes:** **Corresponding author**: Hajime Watanabe, Department of Biotechnology, Graduate School of Engineering, Osaka University, 2-1 Yamadaoka, Suita, Osaka, Japan.

## Abstract

The crustacean *Daphnia magna* is an important model in multi-disciplinary scientific fields such as genetics, evolutionary developmental biology, toxicology, and ecology. Recently, draft genome sequence and transcriptome data became publicly available for this species. Genetic transformation by introduction of plasmid DNA into a genome has been achieved. To further advance *D. magna* functional genomics, identification of a screenable marker gene and generation of its mutant are indispensable. Because *Daphnia* is more closely related to insects among crustaceans, we hypothesized that eye color-related genes can function as a marker gene as used in *Drosophila* genetics. We searched orthologs of *Drosophila* eye pigment transporters White, Scarlet, and Brown in the genome of *D. magna*. Amino acid sequence alignment and phylogenetic analysis suggested that *D. magna* has six *white* and one *scarlet* orthologs, but lacks the *brown* ortholog. Due to a multiplicity of *white* orthologs, we analyzed function of the *scarlet* ortholog, *DapmaSt*, using RNA interference. *DapmaSt* RNAi embryos showed disappearance of black pigments both in the compound eye and in the ocellus, suggesting that *DapmaSt* is necessary for black pigmentation in *Daphnia* eyes. To disrupt *DapmaSt* by using the Crispr/Cas9 system, we co-injected *DapmaSt*-targeting gRNAs with Cas9 mRNAs into eggs and established white-eyed *DapmaSt* mutant lines that lack eye pigments throughout their lifespan. Our results suggest that *DapmaSt* can be used as a transformation marker in *D. magng+a* and the *DapmaSt* mutants would be an important resource for genetic transformation of this species in the future.

## INTRODUCTION

The branchiopod crustacean, *Daphnia magna*, commonly known as the water flea, is a model organism for various scientific fields such as genetics, evolutionary developmental biology, toxicology, and ecology. Its draft genome sequence, together with transcriptome data, is publically available (Orsini *et al.* 2016). Gene manipulation tools such as RNA interference (Kato *et al.* 2011) and genome editing (Nakanishi *et al.* 2014) have been developed. Because *D. magna* is most closely related to insect species among genetic models in arthropods (Schwentner *et al.* 2017), it is suitable for tracing deeper evolutionary roots of developmental programs that have led to the tremendous diversity of arthropod species. To investigate its body plan and sex determination, transformation had been performed using random integration (Kato *et al.* 2012), TALEN- and Crispr/Cas-mediated knockin of plasmid DNA (Nong *et al.* 2017)(Kumagai *et al.* 2017). *Daphnia* has also been long used as a model in ecotoxicology because it is highly sensitive to environmental changes and artificial chemicals (Ebert 2005). To evaluate chemicals with hormone-like activities on this species at the level of gene expression, generation of the biosensor daphniid harboring a reporter gene that responds to ecdysteroids or juvenile hormones has been attempted (Asada *et al.* 2014a)(Nakanishi *et al.* 2016). In these transformation experiments, fluorescent protein genes were used as a visible marker.

Eye color-related genes function as a transformation marker in *Drosophila melanogaster* (Klemenz *et al.* 1987). The fly has three eye pigment transporters: White, Scarlet and Brown, all of which belong to the ATP-binding cassette (ABC) transporter subfamily G (ABCG) harboring one nucleotide-binding domain (NBD) and one transmembrane domain (TMD) (Mount 1987)(Schmitz *et al.* 2001). Each “half ABC transporter” is localized in pigment granule membranes of eye pigment cells (Mackenzie *et al.* 2000) and forms a heterodimer with either of the other two half ABCGs to create a functional transporter. White and Scarlet complex transports a tryptophan-derived precursor, 3-hydroxykynurenine, from a cytosol to a pigment granule, resulting in generation of a brown-colored ommochrome pigment while White and Brown heterodimer transports a guanine-derived precursor that leads to production of a bright red pigment, namely drosopterin (Ewart and Howells 1998). The *white* disruption impairs transport of both pigments and results in change of the compound eye color from red-brown to white. Co-integration of the wild-type *white* with gene-of-interest in its mutant allows us to identify transgene integration events (Klemenz *et al.* 1987). Because eye pigmentation does not need exogenous substrates, and is detectable without special equipment such as a fluorescent microscope, eye color genes would be simpler and more convenient to use than fluorescent protein genes as a transformation marker. However, generation of the mutant line is required.

*Daphnia* has two different types of eyes, a single bilaterally symmetrical compound eye and a single eye or an ocellus at juvenile and adult stages (Güldner and Wolff 1970)(Smirnov 2017). During embryogenesis under laboratory culture at 22 °C, two lateral groups of red ommatidia are first developed at around 36 h post-ovulation (hpo), gradually become close together with turning their color to black at 42 hpo and finally fused along the midline at 48 hpo. Development of an ocellus is also observed at 36 hpo and this eye appears to show black color just after its emergence. The structure of ommatidia has been shown to be similar between crustaceans and insects (Harzsch *et al.* 2007). Presence of *white* and *scarlet* orthologs in a genome of the closely related daphniid *Daphnia pulex* had been reported (Sturm *et al.* 2009). Taken together, it is reasonable to hypothesize that the ABCG orthologs are involved in pigmentation of *Daphnia* eyes. In this study, we annotated orthologs of *white* and *scarlet* in *D. magna*. Knockdown of *scarlet* altered the coloration of compound eye from black to white. We generated white eyed *D. magna* mutant lacking *scarlet* function by using the CRISPR/Cas9 system, which will be used in the development of a transformation marker system in *D. magna*.

## MATERIALS AND METHODS

### *Daphnia* strain and culture conditions

The *D. magna* strain (NIES clone) was obtained from the National Institute of Environmental Studies (NIES; Tsukuba, Japan) and has been cultured under laboratory conditions for many generations. The strain was maintained under the following conditions: 80 neonates (under 24 h) were transferred to 5 L of ADaM medium (Klüttgen *et al.* 1994) and cultured at 22–24 °C under a light/dark photoperiod of 16 h/8 h. The culture medium was changed once after the first week of cultivation. Daphniids were fed once a day with 5.6 × 10^8^ cells/ml *Chlorella* during the first week; after they matured, their offspring were removed once per day and they were fed with 1.12 × 10^9^ cells/ml *Chlorella* daily.

### Bioinformatics analysis of eye pigment transporters

The genomic location of each of the orthologs of eye pigment transporters in insects, *scarlet*, *white* and *brown*, was investigated by tblastn searches using database of the EvidentialGene: *Daphnia magna* Genome (http://arthropods.eugenes.org/EvidentialGene/daphnia/daphnia_magna_new/BLAST/). Amino acid sequences of orthologs from *Daphnia pulex* and *Drosophila melanogaster* (**Table S1**) were obtained from the NCBI database (http://www.ncbi.nlm.nih.gov/) and used as a query. For further confirmation, alignment and phylogenetic trees were constructed using amino acid sequence of each protein. Insect White, Scarlet, and Brown amino acid sequences were obtained from the database (**Table S1**). A multiple alignment was constructed using Clustal W with the following settings: pairwise alignment parameters: gap opening penalty 6.00, gap extension penalty 0.21, identity protein weight matrix; multiple alignment parameters: gap opening penalty 10.00, gap extension penalty 0.24, delay divergent cutoff 30%, gap separation distance 4. Then the phylogenetic tree was constructed using the p-distance algorithm and the neighbor-joining method implemented in MEGA version 7 (Kumar *et al.* 2015). The phylogenetic tree was rooted to *Homo sapiens* ABCG2 family.

### RNA interference of *scarlet*

An siRNA targeting the *scarlet* gene was designed and injected into *Daphnia* eggs according to the method established previously (Kato *et al.* 2011). The sequences of siRNAs were as follows: Scarlet, 5′-GGGUCGCAUUGCUUAUCAA-3′; Control, 5′-GGUUUAAGCCGCCUCACAU-3′ (Asada *et al.* 2014b). Two nucleotides dTdT were added to each 3′ end of the siRNA strand. Briefly, eggs were collected from daphniids, within 2–3 weeks of age, just after ovulation, and placed in ice-chilled M4 medium (Elendt and Bias 1990) containing 80 mM sucrose (M4-sucrose). The siRNA (100 μM) was mixed with 1 mM Lucifer Yellow, which was used as a marker to check injection volume. Then each injected egg was transferred into a well of a 96-well plate filled with 100 μl of M4-sucrose. Injected eggs were then allowed to develop and each injected individual was screened based on eye pigmentation.

### Quantitative real-time PCR

The RNAi embryos were collected at 52 h after injection and homogenized with the beads using the Micro Smash machine MS-100 (TOMY; Tokyo, Japan) in the presence of the Sepasol-RNA I reagent (Nacalai Tesque Inc.; Kyoto, Japan). Total RNA was isolated according to the manufacture’s protocol and followed by phenol-chloroform extraction. The first strand cDNAs were synthesized with the Superscript III Reverse Transcriptase (Invitrogen; Carlsbad, CA, USA) utilizing random primers (Invitrogen) according to the manufacture’s protocol. Quantitative PCR was performed using an Mx3005P real time (RT)-PCR System (Agilent Technologies; CA, USA) with SYBR GreenER qPCR Supermix Universal Kit (Invitrogen) in the presence of a set of primers (*st*-forward 5′-TCTGCGATGAACCAACTACCG-3′ and *st*-reverse 5′-TTTCCGACGAAGGCTGATG-3′). The PCR amplifications were performed in triplicate using the following conditions: 2 min at 50 and 10 min at 95 °C followed by 40 cycles of 15 s at 95°C and 1 min at 60 °C. Gel electrophoresis and melting curve analyses were performed to confirm the correct amplicon size and the absence of the nonspecific band. The target mRNA transcript level was normalized to the transcript level of ribosomal protein L32 (Kato *et al.* 2010). Three biological replicates were used in this experiment.

### Cloning of the *D. magna scarlet* cDNA

Total RNAs were extracted from embryos at 24–54 h after ovulation and cDNAs were synthesized as described above. Based on the cDNA sequence annotated in this study, the gene-specific primers were designed and the partial cDNA fragments were amplified by PCR. The full length *scarlet* cDNA was determined by 5’ and 3’ rapid amplification of cDNA ends (RACE) with a GeneRacer Kit (Invitrogen) and a SMARTer RACE cDNA Amplification Kit (Clontech Laboratories, Inc.; Mountain View, USA). The oligonucleotides sequences for RACE are shown in **Table S2**.

### Knockout of *Scarlet* by CRISPR/Cas9 system

Syntheses of gRNAs and Cas9 mRNAs were performed as described previously (Nakanishi *et al.* 2014). The target site of the *scarlet*-targeting gRNA was 5′-GGTTCACTCGTCGCCTTAATggg-3′ (protospacer adjacent motif shown in lowercase). To generate the gRNA expression vectors, the plasmid pDR274 (Addgene plasmid 42250, (Hwang *et al.* 2013) was digested with *Bsa*I (NEW ENGLAND Biolabs, Connecticut, USA), followed by dephosphorylation with Antartic Phosphatase (NEW ENGLAND Biolabs, Connecticut, USA). Then a pair of *scarlet-*targeting oligonucleotides was annealed and ligated into the linearized pDR274 vector using a ligation mix (TaKaRa Bio, Shiga, Japan). To synthesize gRNAs *in vitro*, gRNA synthesis vectors were digested by *Dra*I (NEW ENGLAND Biolabs, Connecticut, USA) and purified by phenol/chloroform extraction. *Dra*I-digested DNA fragments were used as templates for *in vitro* transcription with mMessage mMachine T7 Kit (Life Technologies, California, USA), followed by column purification with miniQuick Spin RNA columns (Roche diagnostics GmbH, Mannheim, Germany), phenol/chloroform extraction, ethanol precipitation, and dissolution in DNase/RNase-free water (Life Technologies, California, USA).

For synthesis of Cas9 mRNA, a template with the T7 promoter was amplified by PCR from pCS-Dmavas-Cas9 (Nakanishi *et al.* 2014). The amplified PCR fragment was used as a template for *in vitro* transcription with the mMessage mMachine T7 Kit. Poly(A) tails were attached to the capped Cas9 RNAs using the Poly(A) Tailing Kit (Life Technologies, California, USA). The synthesized RNA was then column purified by the miniQuick Spin RNA columns, followed by phenol/chloroform extraction, ethanol precipitation and lastly, dissolution in DNase/Rnase-free water.

gRNA (50 ng/ul) was co-injected with 500 ng/ul Cas9 mRNA into *D. magna* eggs as mentioned above. The eye phenotype of G1 offspring produced by injected G0 mothers was observed under a stereomicroscope. To investigate the Cas9-induced mutations, white-eyed G1 offspring were homogenized in 90 μl of 50 mM NaOH with zirconia beads. The lysate was heated at 95°C for 10 min and then neutralized with 10 μl of 1 mM Tris-HCl (pH 7.5). This crude DNA extract was centrifuged at 12000 rpm for 5 min prior to being used as a template in genomic PCR. The gRNA-targeted genomic regions in *st* locus were amplified by PCR with Ex Taq Hot Start Version (Takara Bio, Shiga, Japan). We used the following primers: *st*-fwd-gDNA 5′-GGTCCCCTTCAAACGAGTC -3′ and *st*-rev-gDNA 5′-GGACATCTGCAAGCCAA-3′. The PCR products were analyzed by polyacrylamide gel electrophoresis and DNA sequencing.

### Reproduction test

Reproduction was assessed with *scarlet* mutants (MT1 and MT2) and wild-type (WT). Twenty-four neonates (<24 h old) were cultured individually in ADaM medium until they produced the third clutch (approx. 21 days at 22–24 °C. They were fed with 5 × 10^6^ cells/ml for 1 week, and thereafter with 1 × 10^7^ cells/ml. The medium was changed and the cumulative number of offspring was counted at each clutch.

### Data Availability

Strains and plasmids are available upon request. The nucleotide sequence of *D. magna scarlet* cDNA is available from the DDBJ database (http://www.ddbj.nig.ac.jp/index-e.html) (accession number LC380914). The authors affirm that all of the other data necessary for confirming the conclusions of the article are present within the article, figures, tables, and supplemental file. Figure S1 shows alignment of orthologs of Scarlet, White, and Brown. Figure S2 include sequences of *scarlet*-targeting siRNA and gRNA in addition to their homology to sequences of *white* orthologs. Table S1 and Table S2 list accession numbers of genes and sequences of oligonucleotides used in this study respectively.

## RESULTS

### Identification of *white* and *scarlet* orthologs in *D. manga* genome

To examine whether orthologs of *white* and *scarlet* exist in *D. magna*, we searched the *D. magna* genome database (Orsini *et al.* 2016). We found six *white* and one *scarlet* genes showing high similarity to *D. pulex*, respectively (**Table 1**). All of the genes code for one NBD and one TMD at N- and C-terminal regions (**Figure 1**). We performed alignment of the seven ABC transporters and found sequence conservation of a Walker A motif (or P-loop), a Walker B motif, a D-loop, a Q-loop, and an H-motif (or Switch region) in a catalytic core domain, in addition to an ABC signature motif in the alpha-helical domain (**Figure 2**, **Figure S1**). The phylogenetic analysis showed that one transporter is grouped into *scarlet* gene and the others into *white* (**Figure 3**). We could not find any brown ortholog in the *D. magna* genome. These results suggest that *D. magna* has a single *scarlet* ortholog and six *white* orthologs.

**Table 1.**
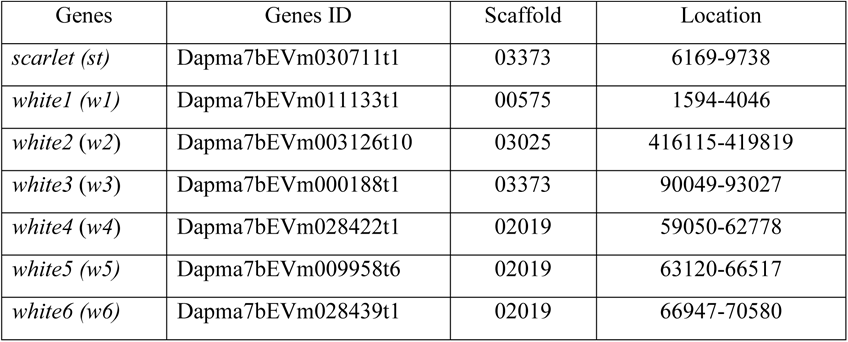
Annotation of *white* and *scarlet* orthologs in *D. magna*.

**Figure 1.**
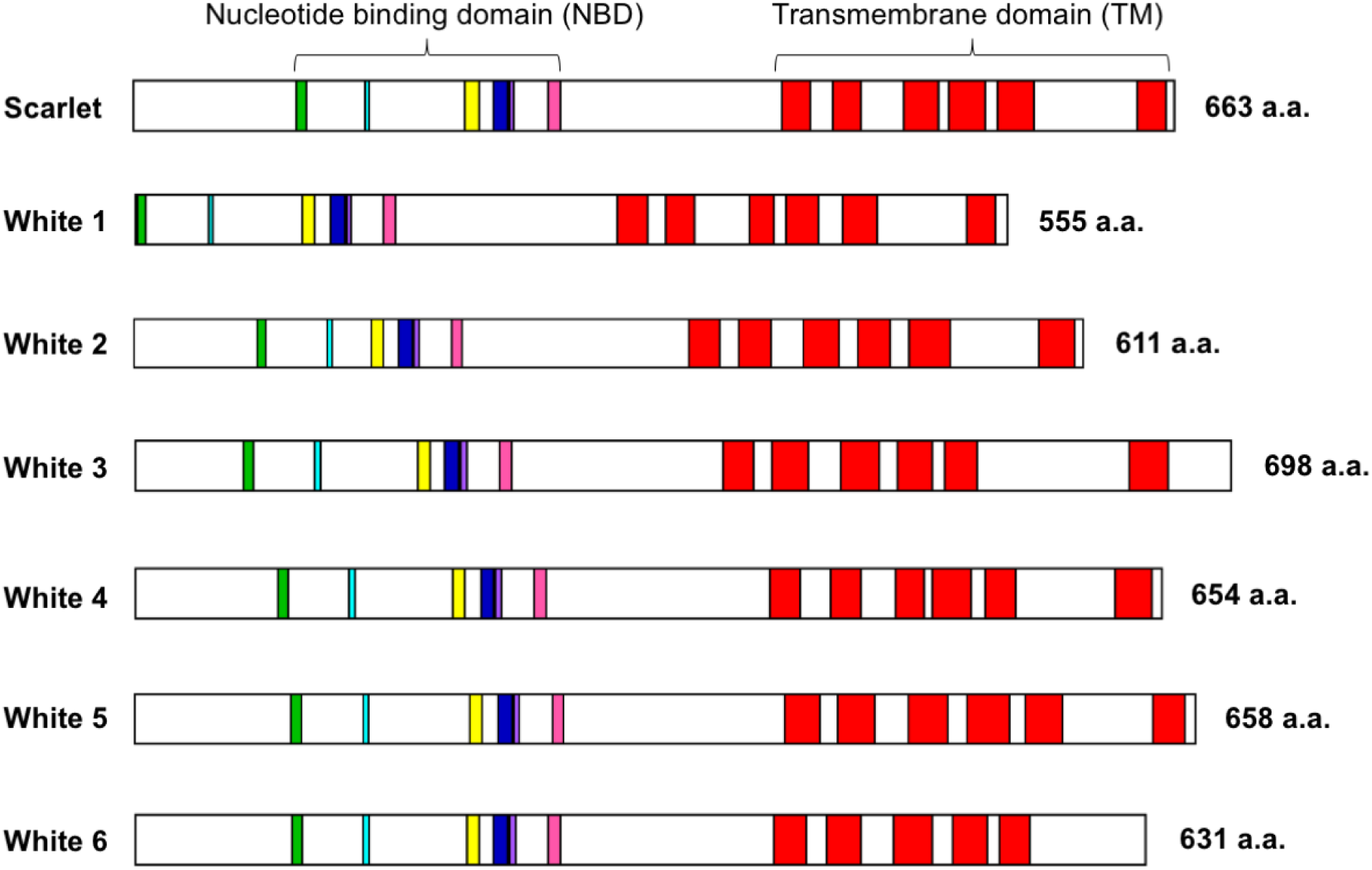
Putative structures of orthologs of eye transporter proteins in Daphnia *magna*. The green, light blue, yellow, dark blue, purple, and pink boxes indicate the Walker A/P-loop, Q-loop, ABC signature motif, Walker B, D-loop, and Switch/H-loop. Red boxes show transmembrane regions.

**Figure 2.**
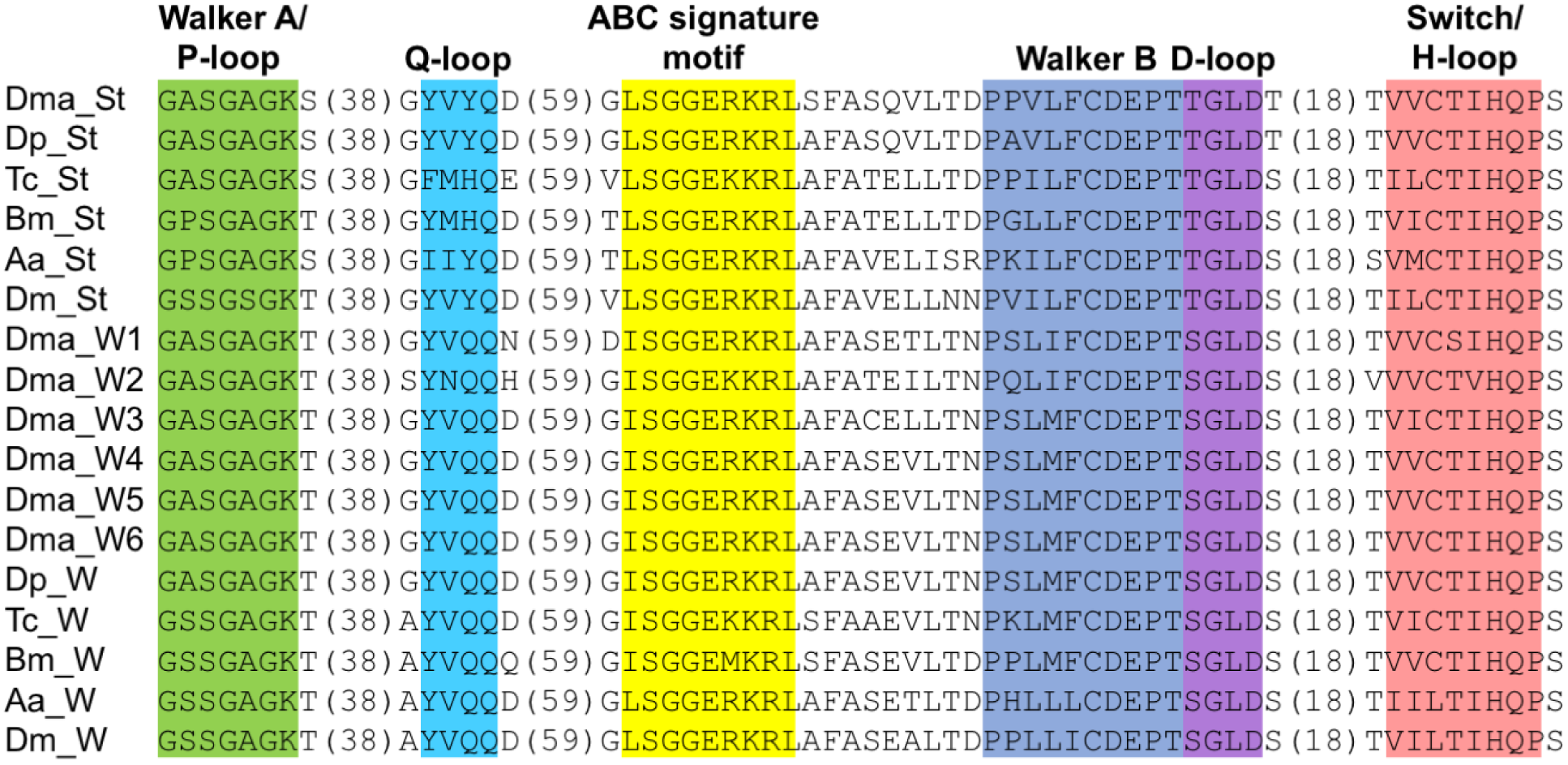
Amino acid alignment of nucleotide binding domains of Scarlet (St), White (W) in *Daphnia magna* (Dma), *Daphnia pulex* (Dp), *Tolibolium castaneum* (Tc), *Bombyx mori* (Bm), *Aedes aegypti* (Aa), and *Drosophila melanogaster* (Dm). The accession number of each protein is shown in **Table S1**.

**Figure 3.**
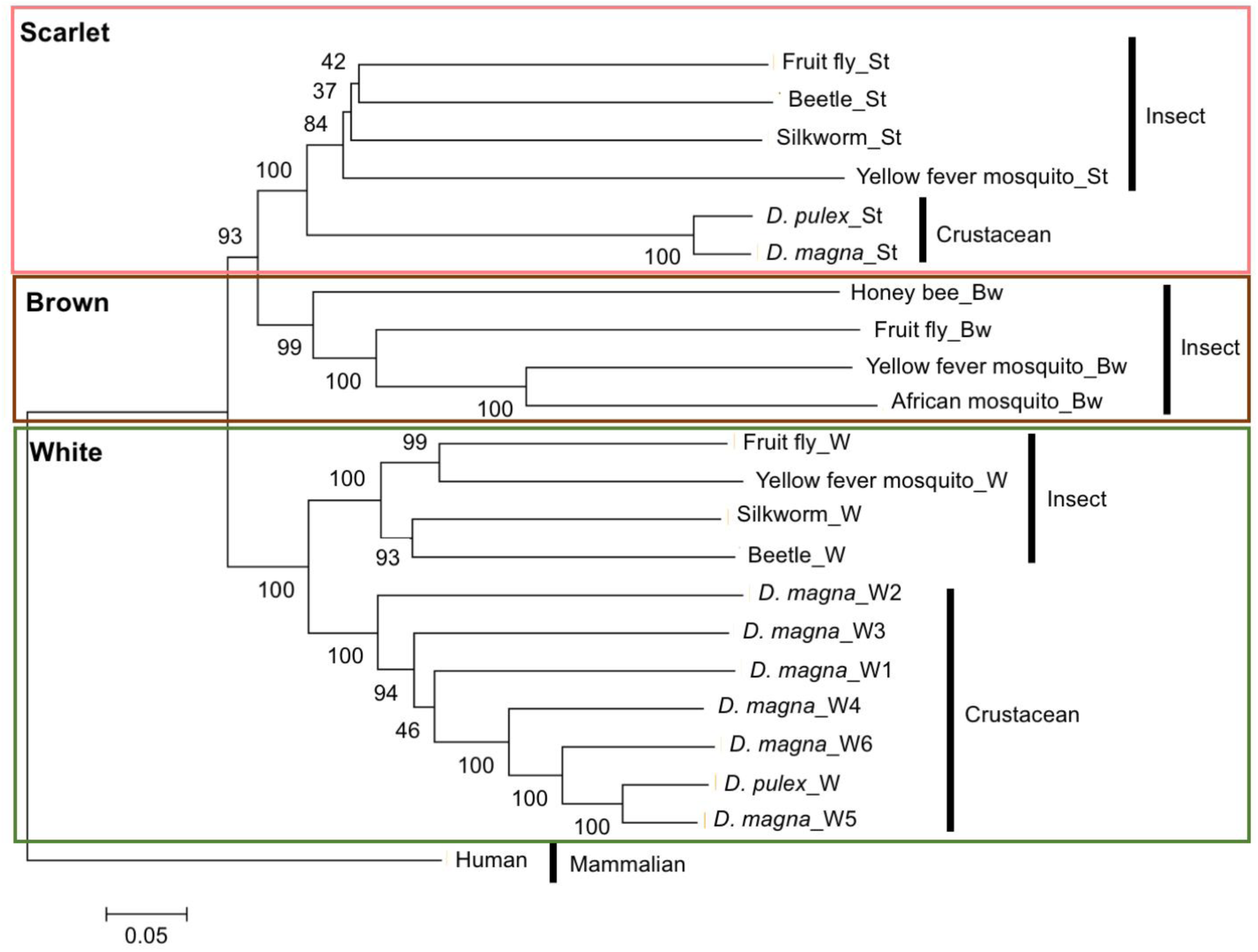
Phylogenetic tree of the amino acid sequences of eye pigment transporters. The percentages of the replicate tree in which the associated taxa clustered together in the bootstrap test (1,000 replicates) are shown next to the branches. The bar indicates branch length and corresponds to the mean number of the differences (*P < 0.05*) per residue along each branch. Evolutionary distances were computed using the p-distance method. The accession number of each protein is shown in **Table S2**.

### *scarlet* is necessary for black pigmentation in eyes

To find a gene involved in eye pigmentation, we focused on the *D. magna scarlet* ortholog (referred as *DapmaSt* hereafter) because multiple *white* genes may lead to gene redundancy that does not produce a clear mutant phenotype. To analyze whether *DapmaSt* is required for black eye pigmentation, we reduced its expression by RNA interference (RNAi). We designed the *DapmaSt*-targeting siRNA in a region that is located downstream of the Switch motif (Figure S1) and is not conserved between DapmaSt and White proteins (Figure S2). We injected it into 29 eggs, and evaluated phenotype of the RNAi at 48 h when a wild-type embryo shows black pigment both in a compound eye and in an ocellus (Figure 4A). All of the injected embryos survived and did not have any pigments in the both types of eyes (Figure 4A) while embryos injected with the control siRNA showed normal eye pigmentation throughout development. The quantity of *DapmaSt* transcripts was decreased to 75% (Figure 4B). These results suggested that *DapmaSt* is necessary for *D. magna* black eye pigmentation.

**Figure 4.**
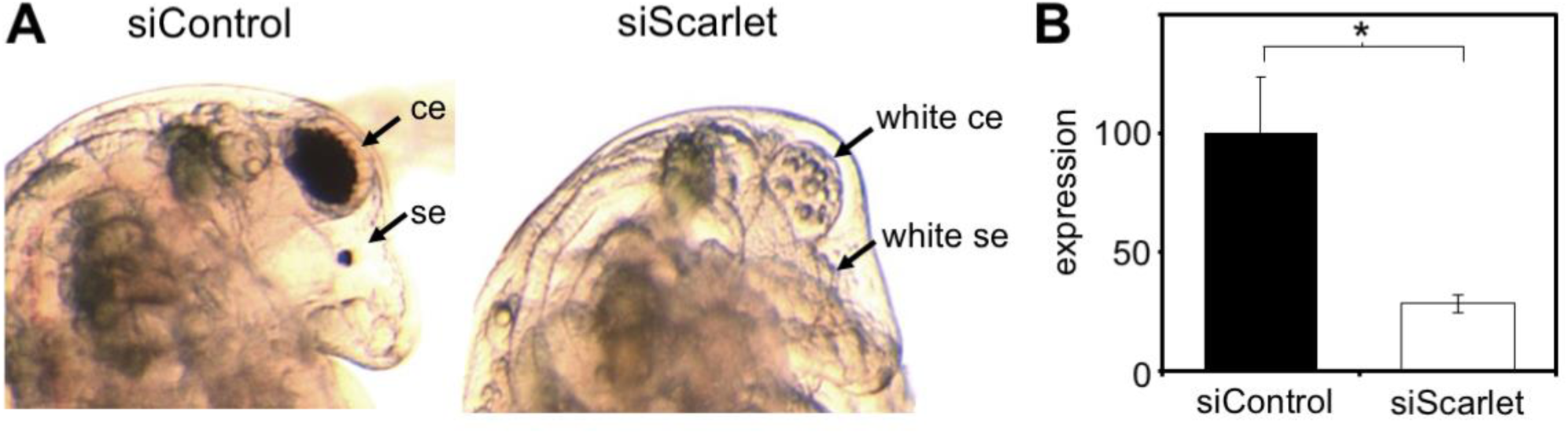
RNA interference of the *scarlet* ortholog in *D. magna*. (A) The typical phenotype of embryos injected with the *Scarlet*-targeting siRNA. The cephalic regions of control siRNA-injected (siControl) and scarlet siRNA-injected (siScarlet) embryos are magnified. The ce and se mean the compound eye and single eye (ocellus). (B) Gene expression profile of *scarlet* in embryos injected with Scarlet siRNAs. Error bars indicate the standard error of the mean (*n* = 3). \**P* < 0.05 (Studet’s t-test).

### Generation of white-eyed *Daphnia* mutants lacking *DapmaSt* gene function

As *DapmaSt* RNAi led to complete loss of eye pigmentation that is similar to a phenotype of *Drosophila white* mutant; the *DapmaSt* mutant might be suitable as a transformation marker in *D. magna*. This prompted us to knockout the *DapmaSt* gene using the CRISPR/Cas9 system. We determined the full-length of the *DapmaSt* ORF sequence by 5′ and 3′ RACE PCRs in the wild-type strain in this study. The sequences were assembled into a transcript that encodes 663 amino acids. Mapping of the sequenced *DapmaSt* cDNA to the genomic sequence indicated that the gene has 13 exons and covers around 3 kb of the genome (Figure 5A).

**Figure 5.**
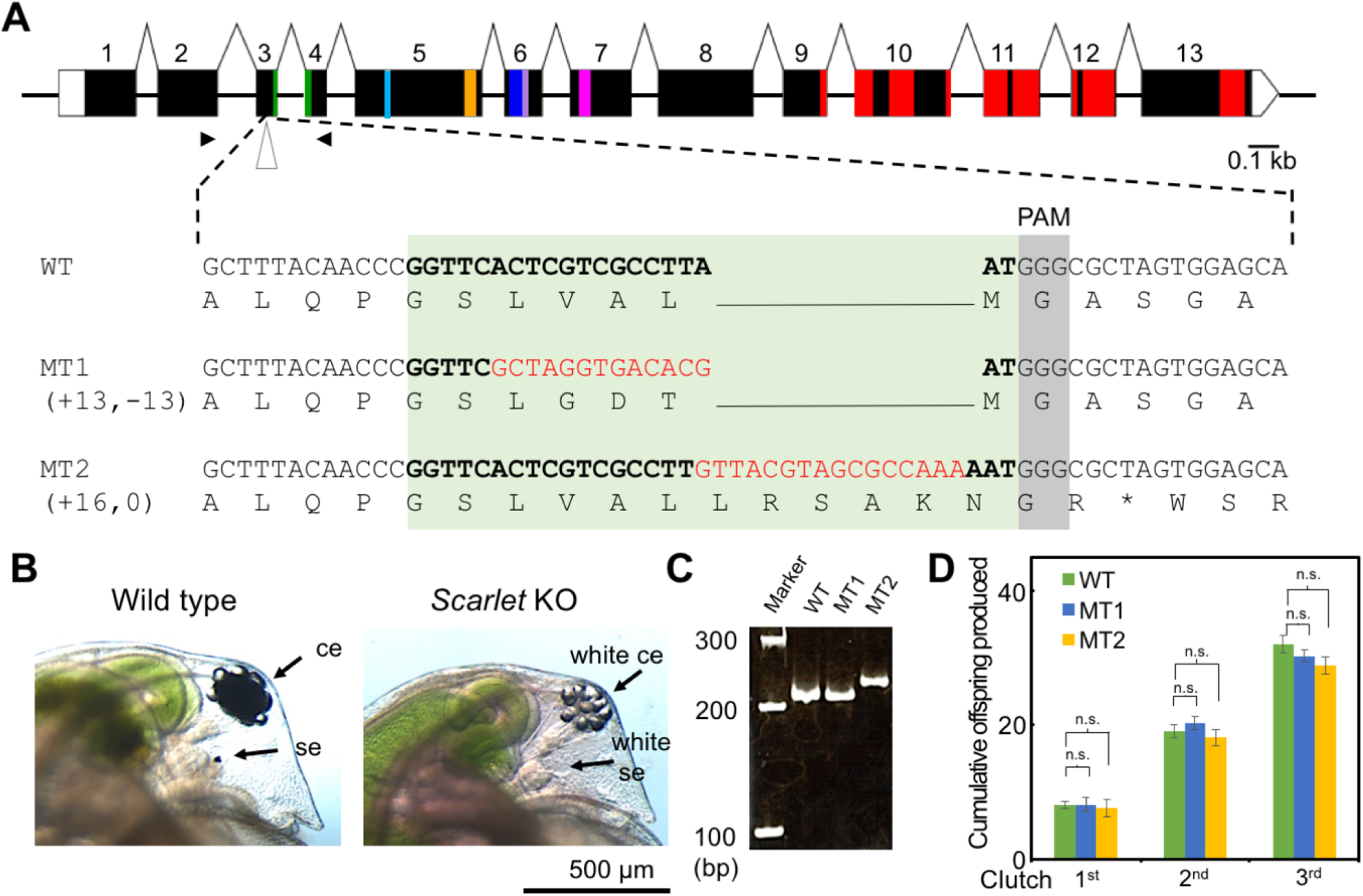
Knockout of the *DapmaSt* in *D. magna*. (A) Schematic gene structure of *DapmaSt* and the partial sequences of a wild-type (WT) and two *DapmaSt* mutants (MT1 and MT2). The green, light blue, yellow, dark blue, purple, and pink boxes indicate the Walker A/P-loop, Q-loop, ABC signature motif, Walker B, D-loop, and Switch/H-loop. Red boxes show transmembrane regions. The target sites for gRNA are indicated by bold letters and insertions are indicated by red letters. The protospacer adjacent motif (PAM) sequence is colored with grey. (B) The phenotype of *DapmaSt* mutant. The cephalic regions of a wild type and the *DapmaSt* mutant are magnified. The ce and se mean the compound eye and single eye (ocellus). (C) Polymerase chain reaction for genotyping of a wild-type (WT) and two *DapmaSt* mutants (MT1 and MT2). The amplified genomic DNA fragments were resolved by agarose gel electrophoresis. (D) Comparison of fecundity between a wild-type (WT) and two *DapmaSt* mutants (MT1 and MT2). The cumulative number of offspring was counted at each clutch. Error bars indicate the standard error of the mean (*n* = 3). n.s. indicates *P* > 0.05 (Student’s t-test).

To disrupt the *DapmaSt* gene specifically on a genome, we designed the gRNA in a region upstream of the Walker A motif (Figure 5A) where its mismatches to the *white* genes are more than 5 bp with PAM (NGG at 3′ end), indicating that off-target to the *white* genes is prevented as reported previously (Jiang *et al.* 2013)(Fu *et al.* 2013). We co-injected 50 ng/μl of the gRNA with 500 ng/μl of Cas9 mRNA. Of 118 injected eggs, 59 survived and became adults. We found two adults that produced G1 progenies with white eyes (founder G0 animals) (Figure 5B) and named progenies from each founder G0 animal as MT1 and MT2 mutants. To investigate the in-del mutations of these mutants at the *DapmaSt* locus, we extracted their genome DNAs, amplified a region around the gRNA-targeted site by PCR, and obtained single PCR fragment for each mutant (Figure 5C). After cloning and sequencing, only one type of in-del mutation for each mutant was identified (Figure 5A). In the MT1 mutant, the indel substituted three amino acids, V74G, A75D, and L76T, indicating that these amino acids are essential for DapmaSt transporter activity. In the MT2, a flameshift mutation occurred and introduced a premature stop codon downstream of the gRNA-target site (Figure 5A). We counted the number of offspring in the first, second, and third clutches from each mutant and confirmed that fecundity of the *DapmaSt* mutants is similar to that of wild-type (Figure 5D). These results demonstrate that the white-eyed *DapmaSt* mutants were established.

## DISCUSSION

Rescue of a visible mutant phenotype, such as loss of eye color, helps our screening of transformants. However, in *D. magna*, any mutant phenotype that is visible and does not affect viability had not been reported yet. In this study, we annotated orthologs of genes that code for eye pigment transporters, White and Scarlet, in insects and analyzed function of the *scarlet* ortholog, *DapmaSt*, by RNA interference and Crispr/Cas9-mediated knockout. We found that *DapmaSt* mutants do not have any pigments in their eyes throughout their lifespan and would become a promising tool for transformation.

Our annotation demonstrated that *D. magna* has six *white* and one *scarlet* orthologs. Sequences of *white* orthologs are widely conserved in arthropod genomes. In a chelicerate (Dermauw *et al.* 2013) and crustaceans, such as copepoda and branchiopoda, including *Daphnia pulex* (Sturm *et al.* 2009)(Jeong *et al.* 2014), multiple *white* orthologs have been identified, but only a single *white* ortholog is present in insects analyzed so far (Dermauw and Van Leeuwen 2014). In contrast, the *scarlet* ortholog has been identified only in insects and daphniids. The *brown* ortholog seems to be insect-specific (Dermauw and Van Leeuwen 2014). Chelicerates branch at the base of the arthropods and insects are nested within crustaceans (Schwentner *et al.* 2017). Among crustaceans, branchiopod crustaceans are more closely related to insects than copepods. Thus, *scarlet* might arise before the common ancestor of insects and branchiopod crustaceans while *brown* appeared to evolve in the insect clade. However, it had not been clarified yet that, except for insects, *white* and *scarlet* orthologs had function in eye pigmentation.

This study revealed that the *D. magna scarlet* ortholog, *DapmaSt*, is involved in pigmentation of a compound eye and an ocellus. *DapmaSt* RNAi caused the white-eyed phenotype. Introduction of mutations into the *scarlet* locus by the Crispr/Cas system dissipated any eye pigments throughout the individual’s lifespan. Unexpectedly, genotyping of two *DapmaSt* mutants, MT1 and MT2, revealed that both mutants have homozygous indel mutations. This might occur due to gene conversion that transfers a mutation on one of the alleles into that on another allele. Alternatively, a large deletion that prevented us from amplification of PCR products was possibly introduced into the target site.

For the purpose of serving as a transgenesis marker, *scarlet* was chosen as a candidate due to only one ortholog found in *D. magna* genome. For transgenic screening, it is better to have a marker gene that can be detected without using any special equipment, such as fluorescence microscope, and its mutant will not affect the development. Previously, we attempted to use the *eyeless* gene; a gene involved in development of *D. magna* eyes, as a transgenesis marker. However, null mutation of this gene is lethal to *D. magna* (Nakanishi *et al.* 2014)(Nakanishi *et al.* 2016), thus making this gene unsuitable as a marker gene. On the other hand, a null mutation of the *scarlet* gene did not affect the development and health of *Daphnia*. This observation is important in deciding what the target marker gene should be.

This work demonstrates identification of *D. magna* orthologs of insect eye pigment transporters and disruption of the *scarlet* gene for generation of mutants with a visible phenotype. The *scarlet* mutant exhibits a white-eyed phenotype, which is similar to the *Drosophila* white mutant. We anticipate that this mutant will be useful for screening of transformants by rescue of the mutant phenotype.

## ACKNOWLEDGEMENTS

This study was partly supported by a grant from the Japan Society from the Promotion of Science (JSPS) KAKENHI (17H01880) to H. W. Y.K. acknowledges the support of the Frontier Research Base for Global Young Researchers, Osaka University, based on Ministry of Education, Culture, Sports, Science, and Technology (MEXT). We would like to thank Editage (www.editage.jp) for English language editing.

